# Contribution of hybridization to the evolution of a derived reproductive strategy in ricefishes

**DOI:** 10.1101/2022.07.05.498713

**Authors:** J. M. Flury, K. Meusemann, S. Martin, L. Hilgers, T. Spanke, A. Böhne, F. Herder, D. Mokodongan, J. Altmüller, D. Wowor, B. Misof, A.W. Nolte, J. Schwarzer

## Abstract

Transitions between reproductive strategies are costly and involve major changes in life history, behaviour and morphology. Nevertheless, in Sulawesi ricefishes, pelvic-brooding evolved from transfer-brooding in two distantly related lineages within in the genera *Adrianichthys* and in *Oryzias*, respectively. Females of pelvic-brooding species carry their eggs attached to their belly until the fry hatches. Despite their phylogenetic distance, both pelvic-brooding lineages share a set of external morphological traits. A recent study found no direct gene-flow between pelvic-brooding lineages suggesting independent evolution of the derived reproductive strategy. It could, however, also be more complex. Pre-existing variation in an admixed population may enable the re-use of genetic variants when subjected to similar external selection pressure, resulting in similar phenotypes.

We thus used a multi-species coalescent (MSC) model and D-statistics to identify gene-tree – species-tree incongruencies, to evaluate the evolution of pelvic-brooding with respect to inter-specific gene-flow not only between pelvic-brooding lineages, but between pelvic-brooding lineages and other Sulawesi ricefish lineages. We found a general network-like evolution in Sulawesi ricefishes and as previously reported, no gene-flow between the pelvic-brooding lineages. Instead, we found hybridization between the ancestor of pelvic-brooding *Oryzias* and the common ancestor of the four *Oryzias* species from Lake Poso, home of the pelvic-brooding *Adrianichthys* lineage. Further, indications of introgression were located within two confidence intervals of quantitative trait loci (QTL) associated with pelvic-brooding in *O. eversi*. We thus hypothesize that a mix of *de novo* mutations and (ancient) standing genetic variation shaped the evolution of pelvic-brooding.

**Significance statement:** The evolution of pelvic-brooding in *Oryzias eversi* (Beloniformes:Adrianichthyidae), was recently described to be independent from another pelvic-brooding ricefish lineage (*Adrianichthys*) from Sulawesi. We confirmed these results, and detected no gene flow between the two distantly related pelvic-brooding lineages. Instead, we found ancient gene flow from another *Orzyias* lineage into the pelvic-brooding *Oryzias* lineage. One of the previously described QTL for pelvic brooding overlaps with a region of high introgression signal. Therefore, we assume that not only *de novo* mutations contributed to the evolution of pelvic-brooding in *Orzyias*, but that introduced ancient genetic variation was likely also recruited for the evolution of this derived brooding strategy.

## Introduction

The evolutionary as well as the genetic basis for the repeatability of evolution exemplified by convergent traits intrigued biologists since the early days (Darwin 1859). Here, similar selective regimes are predicted to result in similar adaptive phenotypes (Schluter & Nagel 1995; Elmer & Meyer 2011). In Sulawesi ricefishes (Beloniformes; Adrianichthyidae), a group of freshwater fishes endemic to the island of Sulawesi, Indonesia, an exceptional reproductive strategy evolved in two distantly related lineages (>15 my divergence time, Hilgers & Schwarzer, 2019; Mokodongan & Yamahira, 2015): This so called “pelvic-brooding” evolved in the genus *Adrianichthys* and in two closely related *Oryzias* species: *O. eversi* and *O. sarasinorum* (Kottelat, 1990; Mokodongan & Yamahira, 2015; Parenti, 2008, Fig. 1). Females of pelvic-brooding ricefish species carry a cluster of eggs attached to their gonoduct for up to three weeks, until the fry hatches (Kottelat 1990). They have elongated pelvic fins and shorter ribs, forming a concavity where the egg cluster is situated (Spanke et al. 2021; Kottelat 1990). In contrast, in the more common and ancestral brooding strategy transfer-brooding (Parenti 2008), females deposit the eggs after several hours (Yamamoto 1975; Wootton & Smith 2014). Changing reproductive strategies entails severe changes in life-history, but also morphological adaptations which in this case are related to prolonged carrying of eggs (Spanke et al. 2021; Parenti 2008; Herder et al. 2012). How such major transitions of reproductive strategies evolve, also on a molecular level, remains unclear.

**Fig 1:**
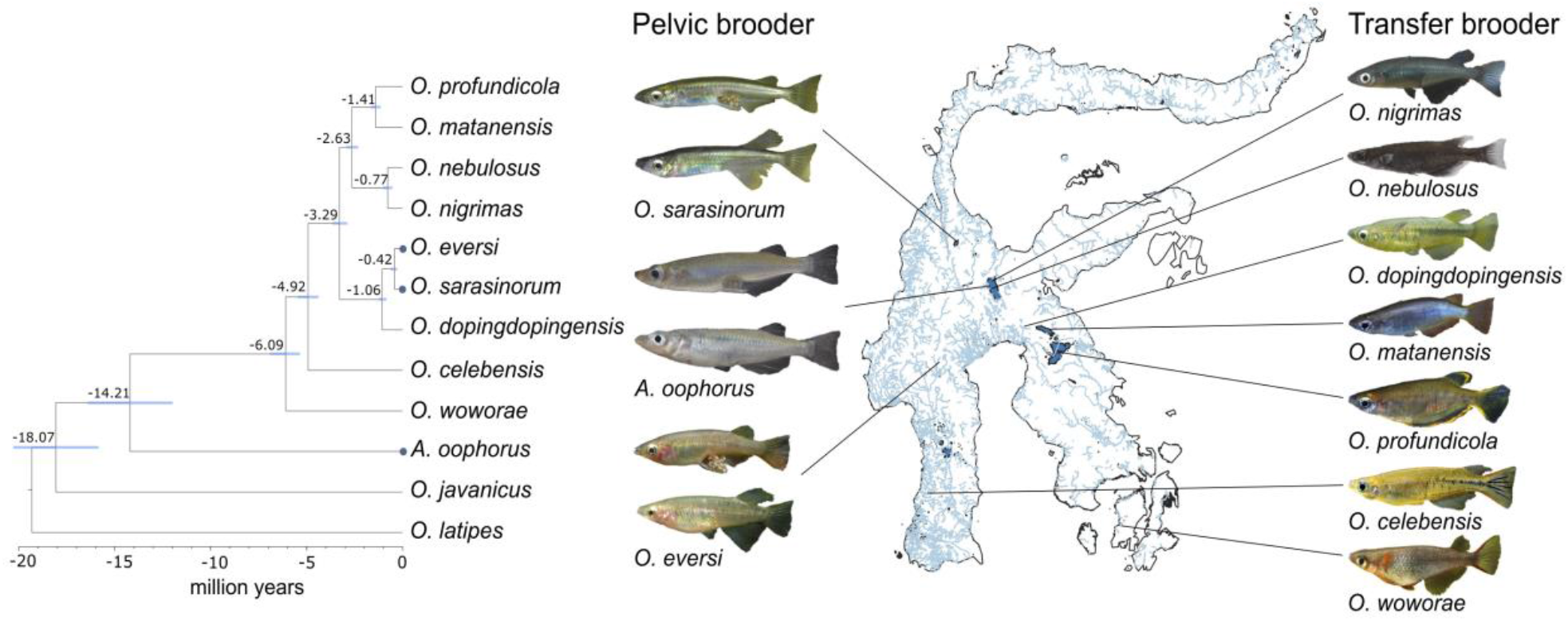
Dated phylogenetic tree adapted from Mokodongan & Yamahira (2015) on the left, pelvic-brooders are marked with blue dots. Map on the right with the distribution of pelvic-brooding (left, male and female showing the pronounced sexual dimorphism) and transfer-brooding species (right, only males). Each clade defined by Mokodongan & Yamahira 2015 for Sulawesi ricefishes, is represented by at least one species. Photos taken by Jan Möhring, Hans Evers and Andreas Wagnitz.

One genetic possibility how the transition from transfer-to pelvic-brooding could be realized are *de novo* mutations. However, they go to fixation rather slowly (Hermisson & Pennings 2005; Barrett & Schluter 2008) and are rarely beneficial (Ohta 1992). Alternatively, the evolution of similar phenotypic traits in different species can also be due to introgression or repeated sorting of shared standing genetic variation. Introgressed variation from one into another population or even species may facilitate the implementation of adaptive variants which are identical by descent (Waters & McCulloch 2021). This so-called adaptive introgression provides fast access to additional genetic variation, if species boundaries are weak (Mallet 2005; Whitney et al. 2006; Arnold 2007; Baack & Rieseberg 2007; Castric et al. 2008; Rieseberg 2009; The Heliconius Genome Consortium et al. 2012; Seehausen 2004). It was proposed that a high level of introgression led to the rapid radiation of *Heliconius* butterflies (Edelman et al. 2019), similar to Darwin’s finches (Lamichhaney et al. 2015) and African Cichlids (Meier et al. 2017). Convergent evolution by repeated sorting (Waters & McCulloch 2021) involves alleles identical by descent, regardless of the potential source of variation, which may be recent (Colosimo et al. 2005; Terekhanova et al. 2014) or rather ancient (Nelson & Cresko 2018; Van Belleghem et al. 2018; van der Valk et al. 2021; Veale & Russello 2017; Brawand et al. 2014).

In the present study, we contrast a convergent evolution scenario of pelvic-brooding with the introgression of genetic variants, by using a phylogenomic approach. A recent study suggested no direct gene-flow between the two pelvic-brooding lineages (Montenegro et al. 2022). However, similar phenotypes may result from recombining pre-existing variation in an admixed population, where diverging species were exposed to similar external selection pressures and genetic variants were re-used (Schluter & Nagel 1995). Gene flow is abundant in Sulawesi ricefishes and has also been detected between pelvic-brooding *Oryzias* species (*O. sarasinorum* and *O. eversi*) (Horoiwa et al. 2021), within Lake Poso *Oryzias* (*O. nigrimas, O. nebulosus, O. orthognathus* and *O. soerotoi*) (Sutra et al. 2019) and the Malili Lake *Oryzias* (*O. matanensis, O. marmoratus* and *O. profundicola*) (Mandagi et al. 2021) which form the sister clades to the pelvic-brooding *Oryzias* (Fig. 1). Thus, gene flow from other sources into the pelvic-brooding lineages harbours the potential to increase genetic variation impacting the evolution of pelvic-brooding. We reconstructed 1907 gene-trees based on single-copy protein-coding orthologous genes of ten Sulawesi ricefishes, five mainland ricefishes and four outgroup species (two poeciliid and two killifish species each). We used these gene-trees as basis for a multi-species coalescence analysis to investigate gene-tree – species-tree incongruencies and to reconstruct a phylogenetic network. We further evaluated introgression on a genome-wide level on published genomes of 17 Sulawesi ricefishes and two mainland ricefish species.

## Results

### Filtering of single-copy orthologous genes

From the 8552 single-copy protein-coding genes derived from the orthologous gene set of Actinopterygii (ID 7898) from OrthoDB, 7331 genes were retained after removal of genes with identified outlier sequences. Randomly similar aligned sections were identified in 1806 genes, hence we masked them (16.5% of all base pairs). According to the TreeShrink analysis, 507 gene trees showed suspiciously long branches and we discarded them from the data set. Phylotreepruner, another method that uses a phylogenetic approach to identify potential wrongly identified orthologs returned no suspicious genes. For final analyses, we kept 1907 orthologous genes (2.415.561 bp) present in all 16 ricefish and four outgroup species.

### Gene-tree – species-tree incongruencies and one moderately supported node in the ML tree

The topology found in the species tree resulting from Astral (Fig. 2A) is highly congruent with the one pulished by Mokodongan and Yamahira (2015) (Fig. 1). In our tree, however, *O. nigrimas* and *O. nebulosus* formed the sister clade to a clade comprising *O. eversi, O. sarasinorum* and *O. dopingdopingensis*. Matching this inconsistency, we found that these clades had many gene-tree – species-tree incongruencies (Fig. 2B). Even though the posterior support values were “1” in each branching event in the species tree from Astral (Fig.S1), quartet scores imply a high degree of uncertainty (Fig. 2C). We observed almost equal quartet frequencies for three splits (split 3, 4 and 7, Fig. 2C in red). At split 3, which indicates the pelvic-brooding *Oryzias* are sister to *O. dopingdopingensis*, one third of all gene trees supported a sistergroup relationship of the pelvic-brooding *Oryzias* lineage with the Lake Poso *Oryzias* and about one fifth supported that *O. dopingdopingensis* is most closely related to the Lake Poso *Oryzias*.

**Fig 2:**
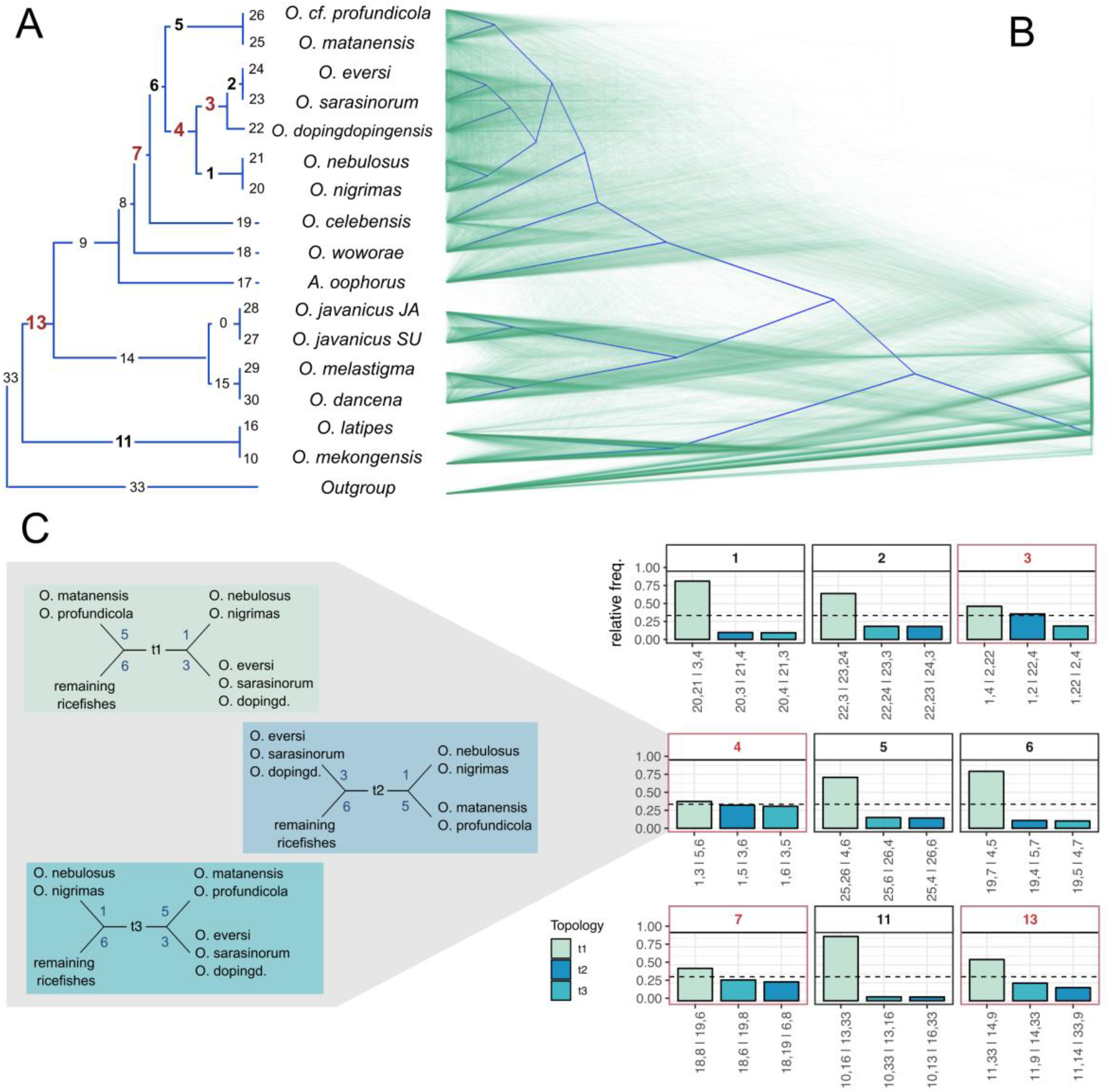
A) species tree from Astral created using Discovista. Numbers in black mean that ∼100% of all gene trees follow the shown topology. Bold numbers refer to quartet scores with >60% of all gene trees following the shown topology. Red numbers show splits with the second most frequent topology found in >25% of all gene trees. B) Densitree based on 1907 single gene trees. Blurry areas indicate gene tree – species tree incongruencies. C) Barplots representing percentage of alternative topologies in corresponding nodes (indicated in red in A). Split 4 is highlighted as example depicting the three different possible topologies.

In split 4 – which was not inferred in the ML tree (Fig. 3), Lake Poso ricefishes were sistergroup to the Malili lake system ricefishes in one third of all gene trees and in one third the Malili lake system ricefishes were sistergroup to the pelvic-brooding species *O. eversi* and *O. sarasinorum* and the transfer brooder *O. dopingdopingensis*. Regarding split 7, in one fourth of all gene trees *O. woworae* is sister to the other Sulawesi *Oryzias* instead of *O. celebensis*, and in another fourth of the gene trees, *O. woworae* and *O. celebensis* were sistergroups and the other Sulawesi *Oryzias* are sister to the rest of the ricefishes. In split 13, comprising also non-Sulawesi *Oryzias*, alternative quartetts occur with about one fourth of all gene trees supporting *O. latipes* and *O. mekongensis* as sistergroup to the Sulawesi ricefishes and one fifth *O. latipes* and *O. mekongensis* are sistergroup to a clade comprising *O. javanicus, O. melastigma* and *O. dancena*.

**Fig 3:**
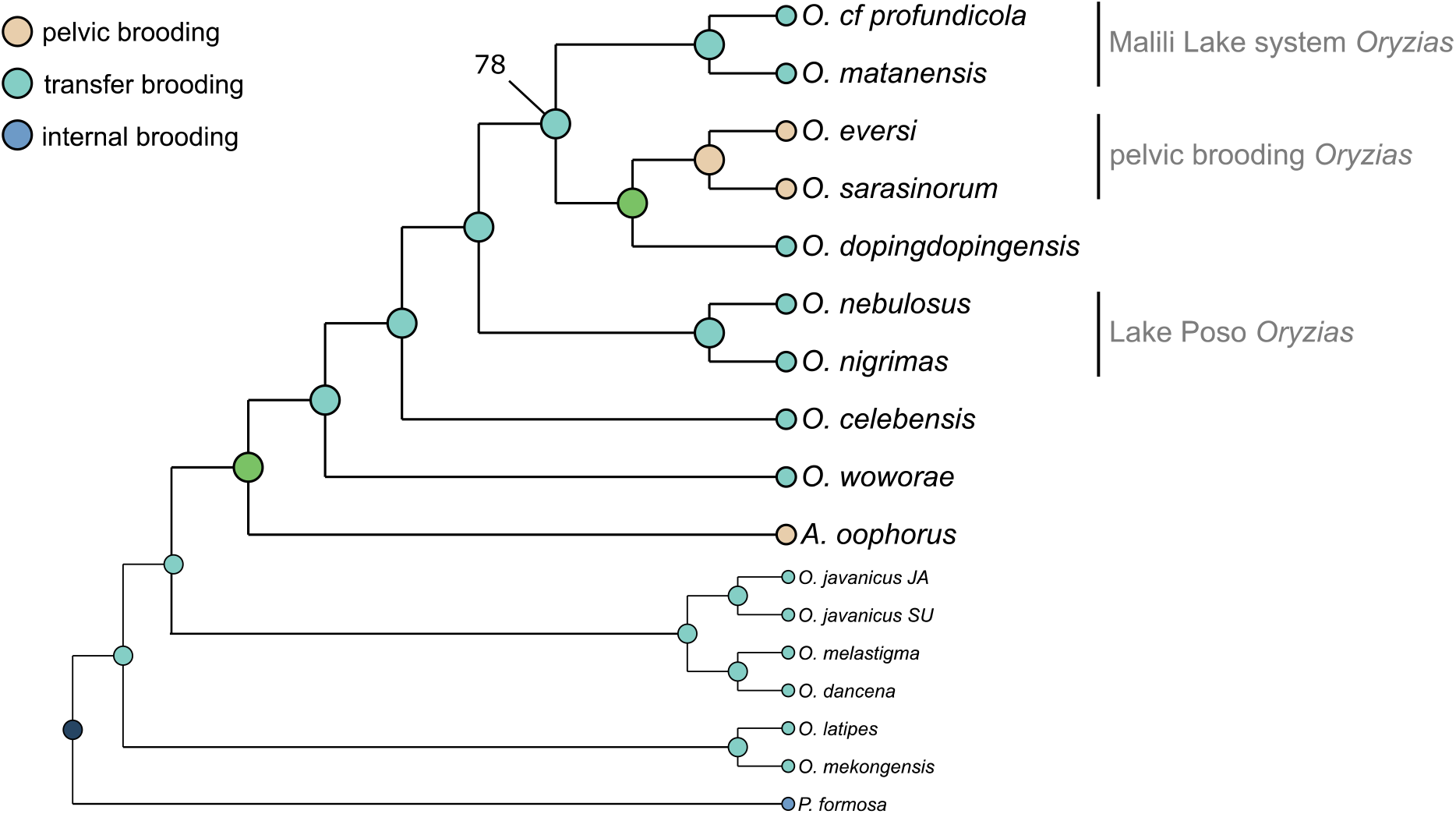
Maximum likelihood tree based on a supermatrix of 1907 concatenated single-copy protein-coding orthologous genes and ancestral state reconstruction. All splits had a bootstrap support of 100, except for the one depicted. Interrelationships between Malili Lake system *Oryzias* and pelvic-brooding *Oryzias* were not well resolved based on our data, indicating that other topologies are also supported (as shown in Figure 2, node 4).

In the ML tree based on the concatenated supermatrix of 1907 orthologous genes, we found a third possible topology (Fig. 3). Here, *O. marmoratus* and *O. matanensis* are sistergroup to *O. eversi, O. sarasinorum* and *O. dopingdopingensis*. All splits obtained maximal bootstrap support, except for the sistergroup relationship between *O. marmoratus, O. matanensis* and *O. eversi, O. sarasinorum* and *O. dopingdopingensis*, which was only moderately supported. The ancestral state reconstruction supported that pelvic-brooding evolved convergently in *Oryzias* and *Adrianichthys* (Fig. 3).

### Detected introgression events in the younger Oryzias clades

We found no introgression among the pelvic-brooding species *A. oophorus, O. eversi* and *O. sarasinorum* (Fig. 4), but between *O. marmoratus* from Mahalona and *O. matanensis* from Lake Matano (Fig. 4). Furthermore, we found a signal of hybridization between *O. sarasinorum* and Lake Poso species *O. nebulosus, O. nigrimas, O. orthognathus* and *O. soerotoi* as well as between *O. eversi* and the same Lake Poso species (Fig. 4). The strongest signals of introgression were located on chromosomes 9, 12_20_13 and 21 (on average >1 window with a *f*_*d*_ and *f*_*dM*_ belonging to the highgest 5% within 30 windows) of the *O. celebensis* reference genome (OCchr) (Fig. S4). Moderatley high signal of introgression was found on OCchr 11, 16 and 24 (on agerage 1 window with a *f*_*d*_ and *f*_*dM*_ belonging to the highgest 5% within 40 windows). For the confidence interval for QCon1 on OCchr 1_17_19 (Tab. S11), a locus associated with the concavity, in 1.46% for *f*_*d*_ and 0.41% for *f*_*dM*_ of the permutations, an equal or higher number of top 1% D-values was found. For the confidence interal of QEgg1, in 9.52% for *f*_*d*_ of the permutations a equal or more negative D-value was found. For QFin on OCchr 24, a proposed QTL was 16 windows away from a window with high introgression single (top 1%). A top 1% *f*_*d*_ value within a range of 15 windows from the proposed location of the QTL was found in 2.73% of the permutations. The other QTLs were on chromosomes with low introgression signal and could not be associated with a high introgression signal (Fig. S4, Tab. S11). The results of gene tree analyses with SNaQ were congruent with results of the D-statistic: all indicated introgression between the Lake Poso *Oryzias* and the pelvic brooders *O. eversi* and *O. sarasinorum* (Fig. 4, Fig. S2). Further, we observed a hybrid edge between the ancestor of *O. matanensis* and *O*. cf. *profundicola*. This was in line with the ambiguous node 4 in the quartet tree of Astral (Fig. 2) and the moderately supported node in the ML tree (Fig. 3). We also observed a hybrid edge between *O. dancena* and the ancestor of *O. celebensis* and the younger *Oryzias* lineages. However, the amount of similarity was rather low with 0.46% (Fig. 4). The genes with the highest D-values were found on OCchr 1_17_19, 8 and 24.

**Fig 4:**
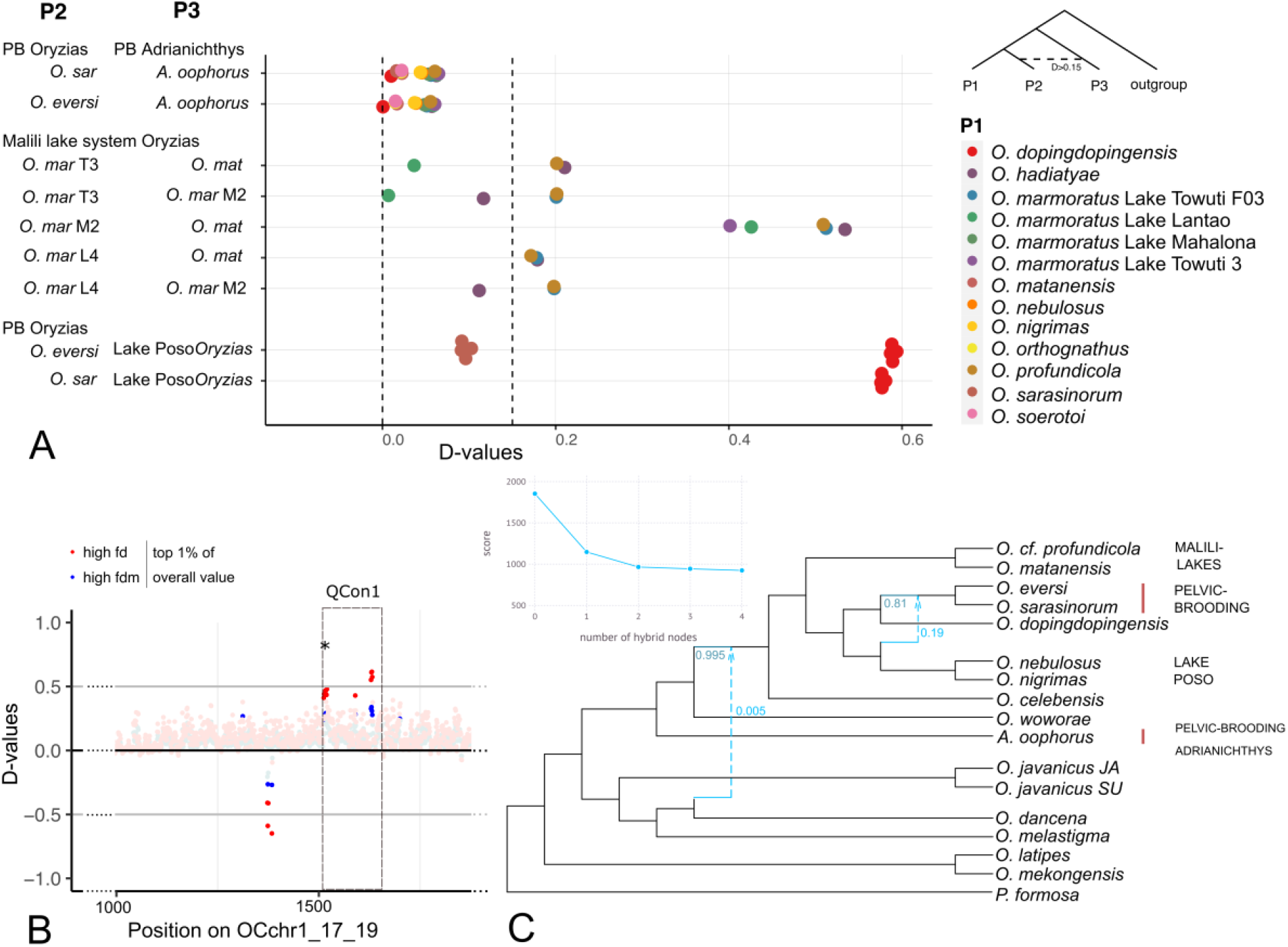
Results from Dsuite and SNaQ. In A) the Dsuite plot, the Lake Poso *Oryzias* refer to *O. nigrimas, O. nebulosus, O. orthognathus* and *O. soerotoi*. For the Malili lake *Oryzias*, O. mar T3 is *O. marmoratus* from Towuti, M2 from Mahalona, L4 from Lantoa. B) shows the region on the *O. celebensis* reference genome chr 1_17_19 with high introgression signal (top 1% of all D-values in color) overlapping with a QTL for the extent of the ventral concavity in *O. eversi*. Both A) Dsuite and C) SNaQ congruently show hybridization between ancestor of Lake Poso *Oryzias* and ancestor of pelvic-brooding *Oryzias*. In the SNaQ tree, another hybrid edge is indicated between *O. dancena* and *O. celebensis* and more recent lineages, though the proportion of genes inherited by this hybrid node from one of its hybrid parents is less than 0.5%.

Using TreeMix (Fig. S3), we identified two introgression events supported with a light migration weight between *A. oophorus* and *O. melastigma* and *O. javanicus* and the most recent common ancestor of the pelvic-brooding *Oryzias* and the Malili Lake system *Oryzias*. With 2 edges (two introgression events), a migration event between the ancestor of the Poso ricefishes and the ancestor of the two *Oryzias* pelvic brooders was suggested, and also had a high migration weight (red arrow). With four edges, the migration event between *O. matanensis* and *O. marmoratus* “Mahalona” was observed. With a fifth edge, another event within the Lake Malili system *Oryzias* was hypothesized.

## Discussion

### No Hybridization between *Adrianichthys* and *Oryzias*, but abundant gene-flow within *Oryzias* from Sulawesi

After analyzing 1907 single-copy protein-coding genes and ∼ 38 mio genomic SNPs, we did not detect any sign of direct gene-flow between the pelvic-brooding lineages, which is in line with previously published results (Montenegro et al. 2022). Slightly different phenotypic expression of pelvic-brooding traits and a differing extent of sexual dimorphism between species of the two pelvic-brooding lineages, *Adrianichthys* and *Oryzias* (Spanke et al. 2021) provide additional support for an independent origin of pelvic-brooding in each lineage. We found, however, gene flow from Lake Poso *Oryzias* into the pelvic-brooding *Oryzias* lineage. Several ancient and recent gene flow events occurred, e.g., locally within lake systems as was shown in the Malili lake system including *O. marmoratus, O. matanensis* and *O. profundicola* (Mandagi et al. 2021). Gene flow over large spatial distances was observed in *O. sarasinorum* and *O. eversi* which nowadays occur 190km apart from each other (Horoiwa et al. 2021). Gene flow over both, large spatial as well as phylogenetic distance, was detected between *O. soerotoi* in Lake Tiu and the ancestor of *O. nebulosus* and *O. orthognathus* in Lake Poso (Horoiwa et al. 2021). All this evidence indicates that ricefish evolution is better described in a network-like evolutionary structure (Fig. 2), rather than by a simple bifurcating tree.

### The quasi-independent origin of a complex reproductive strategy

The QTLs of two traits associated with pelvic-brooding (QCon1 and QFin; Montenegro et al. 2022) lie within regions with high D-values, suggesting that introduced genetic variants from the Lake Poso *Oryzias* might have played a role in the evolution of pelvic-brooding (Fig. S4). However, for two other QTLs the underlying genetic region seemed to have a different genotype compared to the Lake Poso ricefishes and *O. dopingdopingensis* (Qcon2 and QEgg1, expressed by highly negative D-values) and likely evolved *de novo* in the pelvic-brooding *Oryzias*. The introgression event occurred after the splitting of *O. dopingdopingensis* and the pelvic-brooding Oryzias (∼1.4–1.8 mya based on Ansai et al., 2021; Mokodongan & Yamahira, 2015). Hence, a potential gene flow event between the Lake Poso and pelvic-brooding stem lineage dates around the origin of Lake Poso, which is assumed to be not older than 1–2 mya (Rintelen et al. 2004; von Rintelen & Glaubrecht 2006). Intriguingly, all *Adrianichthys* species are endemic to Lake Poso, which would allow an indirect transmission of pelvic-brooding alleles into the stem lineage of the pelvic-brooding *Oryzias*. This assumption is contradicted by the lack of gene flow detected between *Adrianichthys* and any of the *Oryzias* species, though old hybridization events might be hard to detect. Based on our data, we found that a small percentage (0.46%; Fig.S1) of the *O. dancena* genome has introgressed into the ancestor of *O. celebensis* and the younger *Oryzias* lineages hinting towards that older hybridization events occurred between different ricefish lineages. Ancient hybridization events have the potential to serve as resource for adaptive genetic variation. Available ecological niche space allows selection to act upon genetic diversity originating from differently combined old alleles, generating adapted and divergent phenotypes (Seehausen 2013, 2004). For example, in the apple maggot *Rhagoletis pomonella*, introgressed genetic regions were found, deriving from an ancient hybridization with Mexican Altiplano highland fruit flies about 1.6 million years ago. However, the emergence of new species using this old variation to adapt to new hosts happened more recently (Feder et al. 2003). Under this combinatorial view, reassembling old variants into new combinations promotes rapid speciation and adaptive radiation (reviewed in Marques et al. 2019). To further disentangle the genomic background of pelvic-brooding in the two distantly related ricefish lineages, in-depth comparative genomic studies are needed.

### Habitat dependence – why did pelvic-brooding evolve twice?

Irrespective of the genetic background, strong selective pressures must have shaped the evolution from ancestral transfer-brooding to the extended care (of pelvic-brooding), which is presumably costly in terms of increased predation risk and increased energetic costs of the care-giver (Wootton & Smith 2014; Cooke et al. 2006). Especially when assuming convergent evolution the role of the environment was proven to be strong for the adaptive value of replicated trait differentiation (Losos et al. 1998; Gompel & Prud’homme 2009; Arendt & Reznick 2008). Present day macro-habitats of pelvic-brooding ricefishes range from lakes to small karst-ponds (reviewed in Parenti, 2008; e.g. Herder et al., 2012; Mandagi, Mokodongan, Tanaka, & Yamahira, 2018); in Lake Poso, pelvic-brooding *Adrianichthys* mainly occupy open-water habitats, leading to the hypothesis that it evolved in adaptation to the absence of suitable spawning substrates in pelagic habitats (Herder et al. 2012). *Oryzias eversi*, however, was described from a small karst pond where potential spawning substrates are abundant (Herder et al. 2012). Thus, present day habitats make it rather complicated to define a general selective regime that might have favoured the evolution of pelvic-brooding. Furthermore, the paleogeographical history of Sulawesis water bodies is barely known and complicated (Wilson & Moss 1999; Hall 2001), leaving plenty of room for speculations, like presumable ancient connections between waterbodies or the existence of a paleolake in central Sulawesi (Utama et al. 2022). Thus, even an ancestral syntopic distribution of *A. oophorus*, the ancestor of *O. eversi* and *O. sarasinorum* and ancestors of the lake Poso *Oryzias* seems possible. Given the costs and direct fitness effects of switching reproductive strategies and the rather fast convergent evolution of pelvic-brooding in two distinct lineages it appears likely that similar environmental and selection pressures in combination with gene flow fueled its evolution.

## Conclusions

Several gene flow events within Sulawesi ricefishes indicate that the Sulawesi ricefish radiation did not follow a tree-like evolution. In this study, we found no introgression between the two distantly related pelvic-brooding lineages, but detected gene flow into the stem lineage of pelvic-brooding *Oryzias* from Lake Poso *Oryzias*. The presence of elevated D-values within the confidence intervals of QTLs for pelvic fin length and the ventral concavity raise the possibility that genetic variants were recruited for the evolution of pelvic-brooding via hybridziation with a closely related species.

## Material & Methods

### Taxon sampling

Fish were collected in Sulawesi between 2011 and 2013 and some species were bred in the aquarium until they were sacrificed (for details see supplementary Tab. S1). We sequenced the transcriptomes of twelve ricefish species (*Adrianichthys oophorus, Oryzias mekongensis, O. dancena, O*. cf. *profundicola, O. nebulosus, O. javanicus, O. sarasinorum, O. eversi, O. nigrimas, O. woworae, O. celebensis, O. matanensis*) and the genome of *O. dopingdopingensis*. Furthermore, a published genome of *O. javanicus* (GenBank Bioproject: PRJNA505405, Biosample: SAMN10417210) and the official gene sets (OGS, used reference genomes in Tab. S6) of two ricefish species (*O. latipes* and *O. melastigma*) and four outgroup species (*Poecilia formosa, Xiphophorus maculatus, Nothobranchius furzeri* and *Austrofundulus limaneus*) derived from the orthologous gene sets of Actinopterygii (TaxID 7898) from OrthoDB (Tab. S1) were included in our analyses.

### Sequencing and orthology prediction

We extracted RNA from complete fish and prepared non-stranded Truseq mRNA libraries which were sequenced on an Illumina Hiseq2000 at the CCG in Cologne (we provide a shortened version of material and methods here, details about all used methods are in the supplement). We used the Qiagen DNeasy Blood &Tissue Kit to extract DNA of *O. dopingdopingensis*. A genomic short read TruSeq DNA PCR free library was prepared by Macrogen sequencing company; finally 15,698,917,340 bases and 103,966,340 reads were sequenced.

The raw transcriptomic reads were quality-filtered using trim-fast.pl from the PoPoolation pipeline (Kofler et al. 2011) and assembled using Trinity v2.8.4. (Grabherr et al. 2011; Haas et al. 2013). A *de novo* whole-genome assembly of *O. dopingdopingensis* short reads was generated according to Böhne et al. (2019) and Malmstrøm, Matschiner, Tørresen, Jakobsen, & Jentoft (2017) and annotated using gene predictions of *O. latipes* and *O. melastigma*, a protein set from *O. javanicus* and a set of orthologous genes generated in this study. Busco completeness scores of the transcriptomes ranged from 60.7% to 80.9% (Actinopterygii Odb9) (Tab. S3) and the annotated genome assembly had 97% complete Busco completeness score.

We created the reference ortholog set using the orthologous gene set of Actinopterygii (TaxID 7898) downloaded from OrthoDB (Waterhouse et al. 2013). We used Orthograph v0.7.1 (Petersen et al. 2017) to find orthologous genes in our transcriptome assemblies and the genome assembly. The orthologous genes were aligned using MAFFT v7.221 with the L-INS-I algorithm (Katoh & Standley 2013) on the amino-acid level and outliers in genes were identified and the respective genes removed according to Misof et al. (2014). We created nucleotide alignments with the amino acid alignments as a control with a modified version of Pal2Nal v14 (Suyama et al. 2006; Misof et al. 2014) and masked ambiguously aligned regions (maximum number of pairwise sequence comparisons for each MSA (option -r), and the -e option for gap-rich data sets, leaving remaining settings to defaults) and subsequently removed all MSA sections, which were indicated as randomly similar aligned with ALISCORE v2.0 and ALICUT v2.3 (Misof & Misof 2009; Kück et al. 2010; Misof et al. 2014).

### Phylogenomic analyses

We used IQ-TREE v1.6.12 to create trees for every gene separately (Nguyen et al. 2014) (for details on the methods described in this paragraph, see supplement). For each gene, a substitution model was estimated (Tab. S9). We used Astral v5.7.3 to run a multi-species coalescent model (Zhang et al. 2018). The gene trees were tested with Pyhlotreepruner (Kocot et al. 2013) and TreeShrink (Mai & Mirarab 2018). Phylotreepruner checks for paralogous sequences and TreeShrink for extremely long branches. The results of Astral were displayed using DiscoVista (Sayyari et al. 2018). The gene trees were made ultrametric using the function “chronos” in R v4.1.2 (package “ape” v5.6-2 (Paradis & Schliep 2019) to use them for Densitree2.01 (Bouckaert & Heled 2014).

The genes were ordered according to their position on the reference genome of *O. latipes* (RefSeq GCF_002234675.1, Bioprject PRJNA325079) and in this order concatenated using FASconCAT-G (Kück & Longo 2014). The resulting supermatrix was partitioned and for each partition a suitable model was searched in IQ-TREE, taking codon position into account. We ran 20 single tree searches and 50 non-parametric, slow bootstrap replicates. Finally, we did an ancestral state reconstruction in RASP4.2 (Yu et al. 2020). We assigned a state of pelvic-brooding to *O. eversi, O. sarasinorum* and *A. oophorus*, a state of transfer brooding to the other ricefishes and a state of live-bearing to the outgroup *poecilia formosa*. We used the ML tree as a basis and ran the S-DIVA model.

### Analyses of gene-flow pattern based on genome-wide SNPs and orthologous genes

We downloaded whole genome data from GenBank of 26 individuals of 19 ricefish species including four specimen of *O. marmoratus*, three *O. wolasi* specimen, and four *O. woworae* specimen from different localities (Bioproject PRJDB10385), *O. javanicus* (Bioproject PRJNA505405, accession number SRR8467745) and *O. melastigma* (Bioproject PRJNA556761, accession number SRR12442554) and mapped them on the *O. celebensis* reference genome from Genbank (GCA_014656515.1, Bioproject PRJDB10371, Ansai et al., 2021) (methods used in this paragraph are described in detail in the supplement). Filtering resulted in 38183142 SNPs and we created a vcf file which we used for Dsuite v0.4r41, a program to calculate D-statistics based on a vcf file (Malinsky et al. 2021). To locate regions of elevated D-values, we used DInvestigate, a follow-up analysis included in Dsuite. It calculates different D-statistics for windows across the genome. We left window size and steps at default. We used *f*_*d*_ and *f*_*dM*_, two statistics specifically designed to detect introgressed loci. Where *f*_*d*_ is distributed on the interval [-Infinity, 1] (Martin et al. 2015), *f*_*dM*_ is distributed on the inveral [-1,1] under the null hypothesis of no introgression is symmetrically distributed around zero (Malinsky et al. 2015). To track down chromosomes with the strongest signal of introgression, windows with top 5% (Fig. S4) values for genome-wide *f*_*d*_ and *f*_*dM*_ were counted and corrected for chromosome length. Additionally, we checked whether windows with the top 5% and top 1% of genome-wide *f*_*d*_ and *f*_*dM*_ values fall within the confidence intervals of QTLs associated with pelvic-brooding (Fig. S5). To test if D-values were significantly higher (if positive) or lower (if negative) than expected by chance in the QTL intervals, we distributed the windows with highest D-values (top 1%) randomly over the whole genome 10’000 times and checked, how often we find the same or higher D-values compared to the real QTL intervals. Further we used Treemix v1.13, a graph-model based program, to asses hybrid edges based on the SNP dataset with all missing data removed (Pickrell & Pritchard 2012). To check for signatures of hybridization within the orthologous genes, we used SNaQ which additionally takes different evolutionary rates of branches into account (Solís-Lemus et al. 2017; Solís-Lemus & Ané 2016). We used HybridCheck to find elevated D-values in our orthologous gene set (Ward & van Oosterhout 2016). The regions of elevated D-values were compared to the quantitative trait loci (QTLs) associated with pelvic fin length, the extent of the concavity and duration of egg carrying found in (Montenegro et al. 2022).

## Acknowledgements

We thank Rob Waterhouse for the advice on using OrthoDB. Further, we are thankful for the help of Alexandros Vasilikopoulos with compiling the ortholog set. We thank Michael Matschiner and Thore Koppetsch for their insights in the pitfalls of different evolutionary rates in hybridization analysis and their recommendations. We are very grateful for the permission to use beautiful photos taken by Jan Möhring, Andreas Wagnitz and Hans Evers for this publication. Calculations for the genome assembly were performed at sciCORE (http://scicore.unibas.ch/) scientific computing center at University of Basel. Further, we used the annotation scripts from the Sigenae platform for the genome annotation. This work was supported by the Leibniz Association, grant P91/ 2016.

## Data Availability Statement

### Genetic data

Raw sequence reads are deposited in the SRA (will follow after acceptance). Genome annotation data is available on DataDryad (will follow after acceptance).

### Sample metadata

Related metadata can be found in supplementary material (will follow after acceptance).

## Benefit-Sharing Statement

Benefits Generated: A research collaboration was developed with scientists from the countries providing genetic samples, all collaborators are included as co-authors, the results of research have been shared with the provider communities and the broader scientific community (see above). More broadly, our group is committed to international scientific partnerships, as well as share knowledge about the establishing and maintaining of scientific collections.

## Author contributions

JMF, JS, KM designed research. AWN and FH were leading the field expeditions in Sulawesi where the samples were collected. AWN provided the transcriptome sequences. LH did the transcriptome assemblies. JMF and KM performed research and analyzed data. AB did the genome assembly. SM annotated the genome and supported JMF with the NCBI submission process. JMF and JS wrote initial draft of the manuscript. All authors contributed to writing the final manuscript.

## Supplementary figures

**Fig S2:**
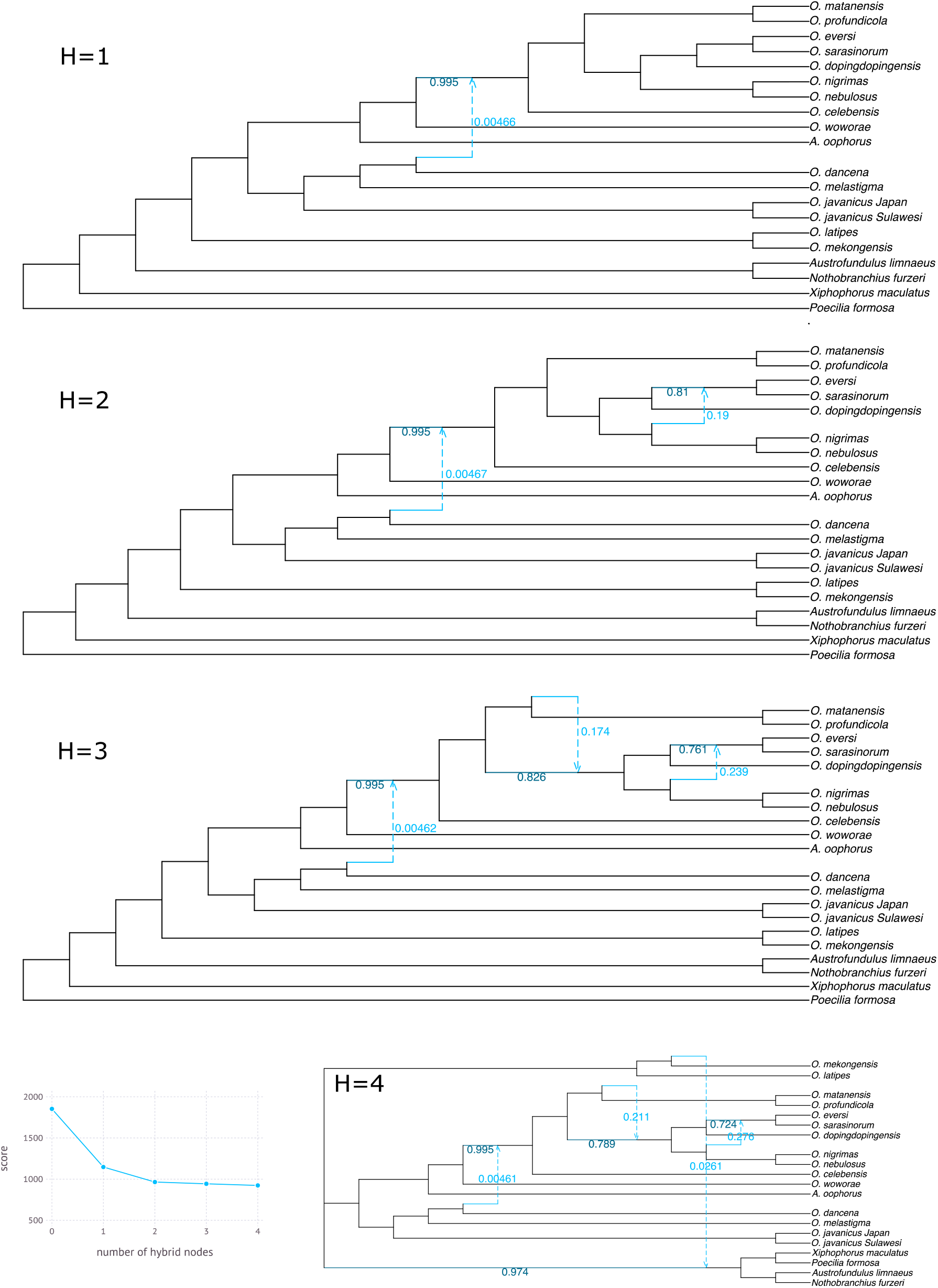
Result from SNaQ for 1-4 hybridization events. Hybridization between *O. mekongensis* and outgroup rather unlikely and probably false positive result.

**Fig S3:**
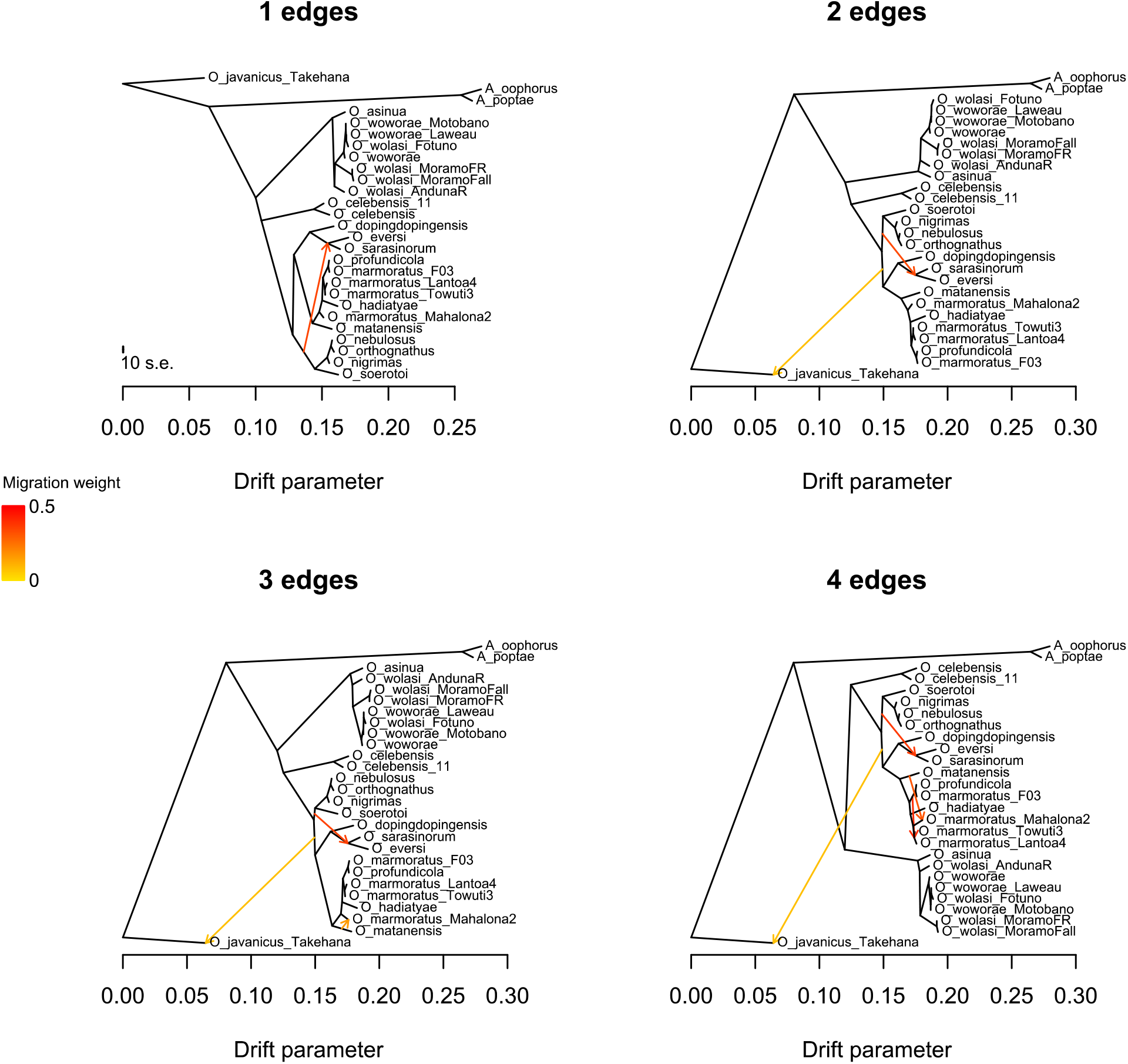
Results from TreeMix: same introgression events were found as in the Dsuite analysis, 1) between the *Oryzias* pelvic brooders and the *Oryzias* Lake Poso and 2) between *O. matanensis* and *O. marmoratus* Mahalona.

**Fig S4:**
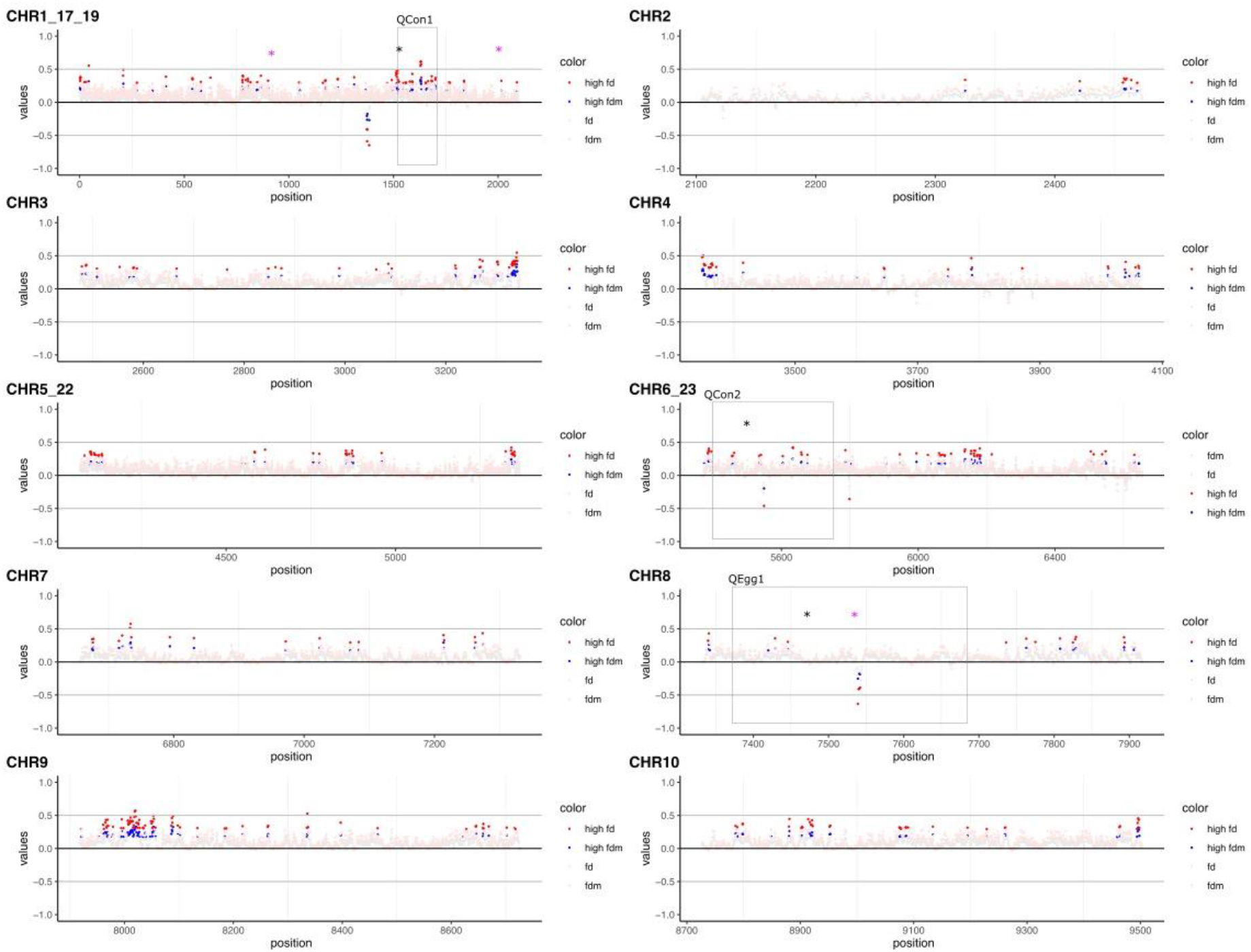
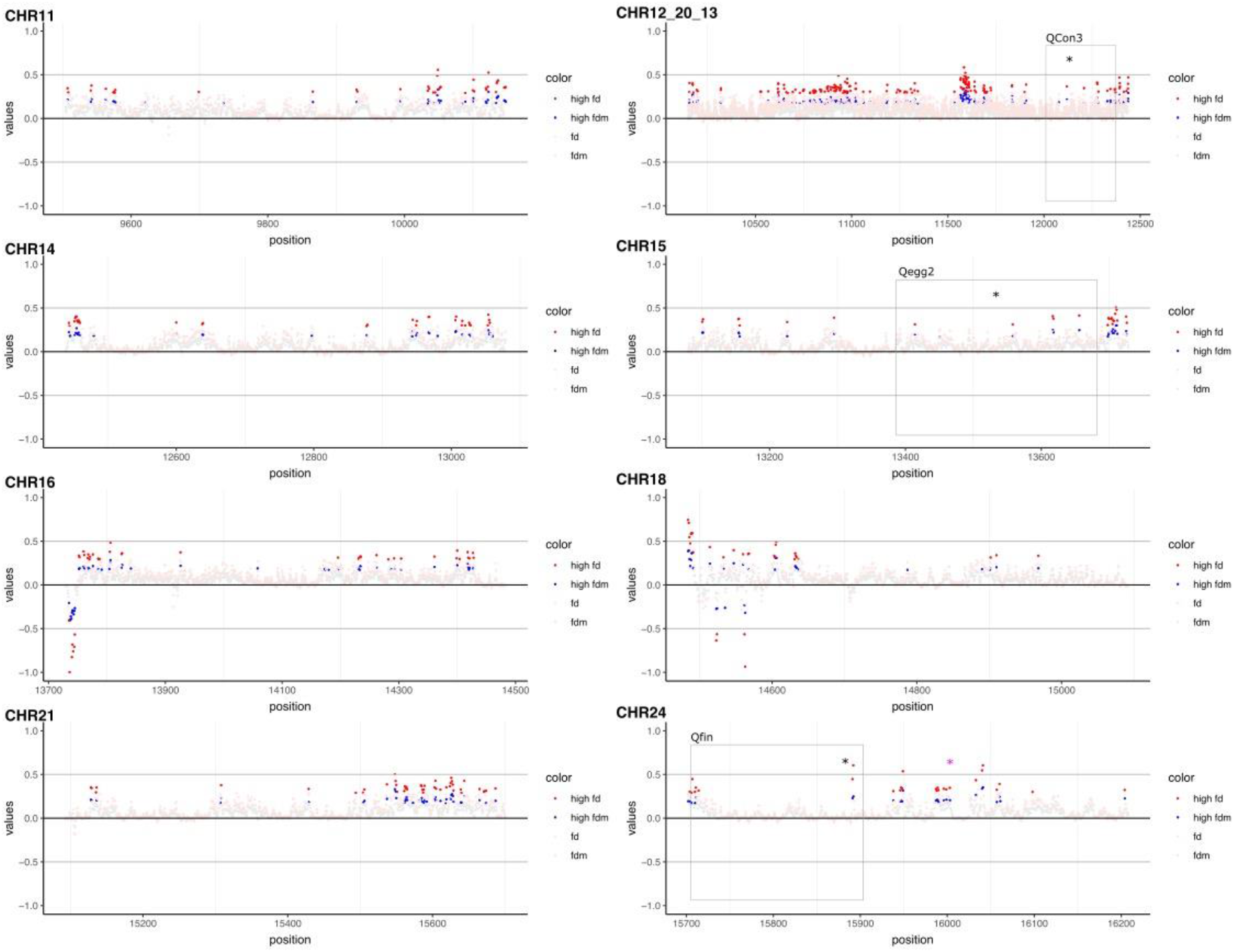
Results from DInvestigate: On the y-axis are the D-values for the sliding-windows (size 50 kbp, red top 5% values of *f*_*d*_, blue top 5% values of *f*_*dM*_). On the x-axis are the positions on the respective *O. celebensis* reference genome, naming follows the synteny with the *O. latipes* reference genom (how positions refer to base pairs can be read in Table S11). Grey boxes mark the confidence intervals of the QTLs found in Montenegro et al., 2022, black stars the QTLs (QCon1-3, QEgg1-2, QFin). Pink stars mark the position of the genes with elevated D-values found in HybridCheck.

**Fig S5:**
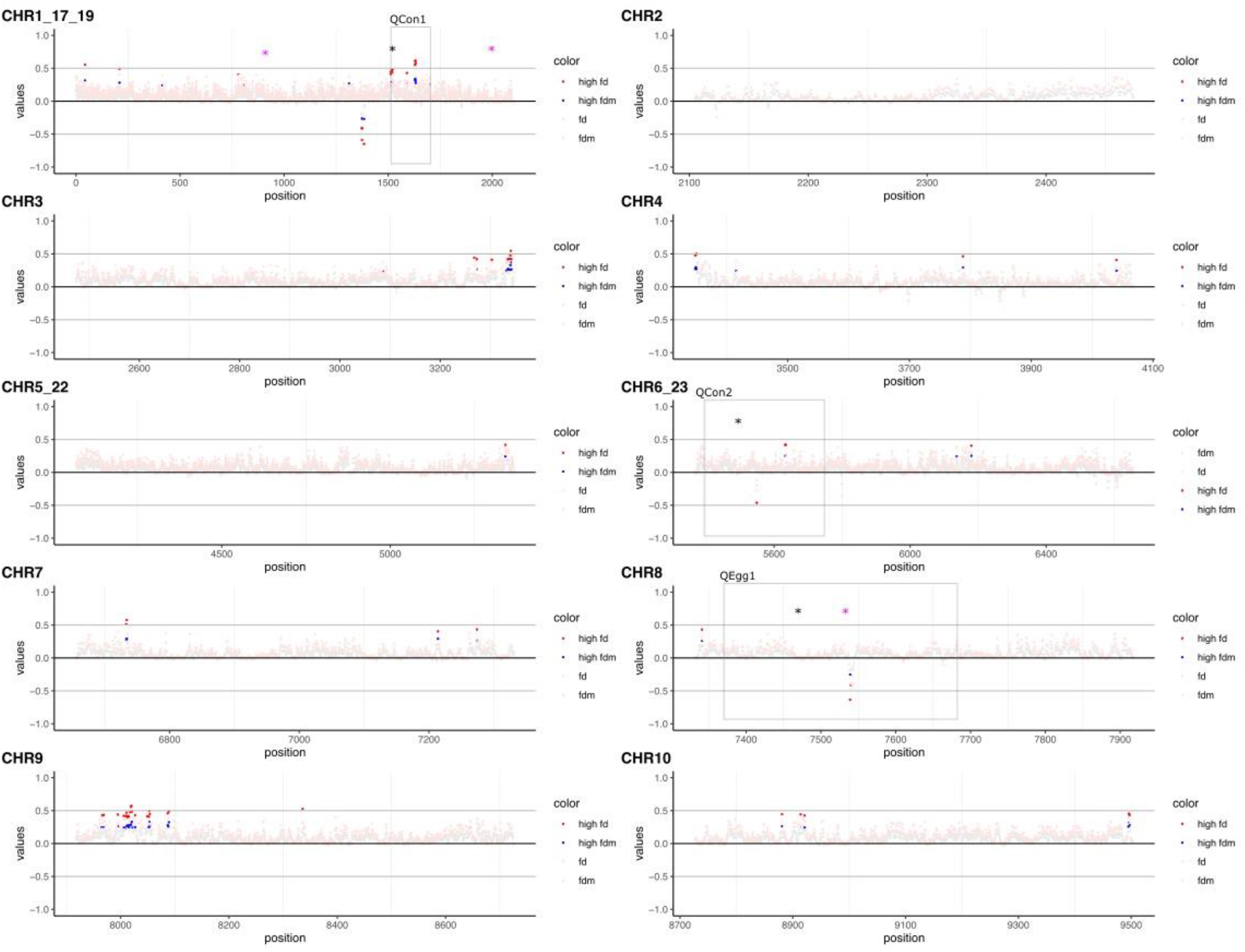
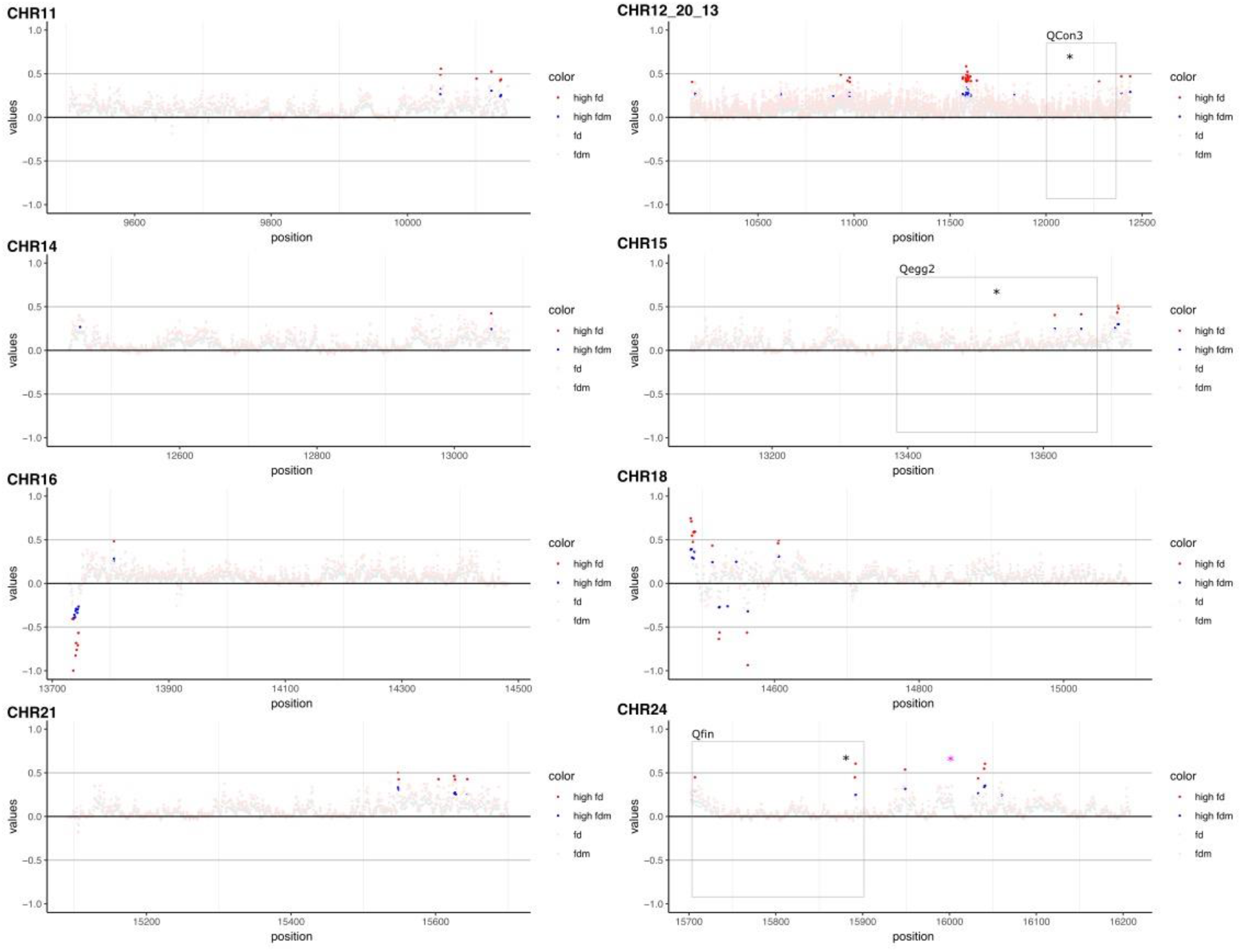
same figure as S4, but with only top 1% of D-values in red and blue. Sliding-windows have a size of 50 kbp, in red top 1% values of *f*_*d*_, in blue top 1% values of *f*_*dM*_). On the x-axis are the positions on the respective *O. celebensis* reference genome, naming follows the synteny with the *O. latipes* reference genome (how positions refer to base pairs can be read in Table S11). Grey boxes mark the confidence intervals of the QTLs found in Montenegro et al., 2022, black stars the QTLs (QCon1-3, QEgg1-2, QFin). Pink stars mark the position of the genes with elevated D-values found in HybridCheck.

